# A protein-based biosensor for detecting calcium by magnetic resonance imaging

**DOI:** 10.1101/2021.02.04.429691

**Authors:** Harun F. Ozbakir, Austin D.C. Miller, Kiara B. Fishman, André F. Martins, Tod E. Kippin, Arnab Mukherjee

**Author notes:** Correspondence should be addressed to A.M.

## Abstract

Calcium-responsive contrast agents for magnetic resonance imaging (MRI) offer an attractive approach to noninvasively image neural activity with wide coverage in deep brain regions. However, current MRI sensors for calcium are based on synthetic architectures fundamentally incompatible with genetic technologies for *in vivo* delivery and targeting. Here, we present a protein-based MRI sensor for calcium, derived from a calcium-binding protein known as calprotectin. Calcium-binding causes calprotectin to sequester manganese. We demonstrate that this mechanism allows calprotectin to alter T_1_ and T_2_ weighted contrast in response to biologically relevant calcium concentrations. Corresponding changes in relaxation times are comparable to synthetic calcium sensors and exceed those of previous protein-based MRI sensors for other neurochemical targets. The biological applicability of calprotectin was established by detecting calcium in lysates prepared from a neuronal cell line. Calprotectin thus represents a promising path towards imaging neural activity by combining the benefits of MRI and protein sensors.

## INTRODUCTION

In the nervous system, calcium ions give rise to intracellular signals that modulate a wide range of biological functions including neural activity, gene expression, synaptic communication, and apoptosis ^[1]^. Consequently, experimental imaging of calcium in living cells and organisms is a cornerstone technology for obtaining detailed, time-lapse information on neural signaling mechanisms. High-resolution calcium imaging typically relies on fluorescent dyes and genetically encoded calcium indicators (GECIs) ^[2]^, which in conjunction with multiphoton microscopy, can detect activity in single cells up to depths of 1 mm in intact tissue ^[3]^. However, the imaging volume accessible by optical methods encompasses only a small fraction of the brain in most vertebrates ^[4]^. Calcium signals can also be recorded from deeper tissues using endoscopes and gradient index lenses, but these techniques cover a limited field of view and require invasive brain surgery to embed the imaging hardware in in brain tissue ^[5]^. Among noninvasive modalities, magnetic resonance imaging (MRI) is unrivaled in its ability to access large volumes of intact tissue located at any arbitrary depth ^[6]^. Thus, MRI-based calcium sensors have the potential to uniquely complement optical indicators by enabling wide-field imaging of biological processes in deep tissues noninvasively. This vision has motivated the development of a versatile collection of small-molecule paramagnetic complexes, fluorinated agents, and superparamagnetic iron oxide crystals that have evolved over the past two decades for imaging calcium by various MRI mechanisms including longitudinal (T_1_) ^[7, 8]^and transverse (T_2_) relaxation ^[9, 10]^, chemical exchange saturation transfer ^[11]^, and direct detection of ^19^F spins ^[12]^. Regardless of the specific contrast mechanism, all reported MRI probes for calcium are synthetic molecules ^[7, 8, 9, 11, 12, 13, 14]^, which makes them incompatible with genetic technologies for *in vivo* delivery, long-term expression, and cell type specific targeting – key aspects that underpin the prolific success and widespread adoption of calcium sensors derived from the green fluorescent protein (GFP) ^[15]^. While genetically encodable iron-containing enzymes and protein nanoparticles have been used to develop MRI sensors for functional imaging of neurotransmitters and kinase activity ^[16, 17, 18, 19]^, no protein-based MRI probe has been reported for calcium. To this end, here we describe the development of the first biomolecular MRI reporter for calcium, based on a novel manganese metalloprotein, and demonstrate its utility for imaging calcium concentrations relevant to intracellular signaling.

## RESULTS

Our sensor is based on calprotectin, an antimicrobial protein that is released by neutrophils to chelate essential transition metals (including paramagnetic Mn^2+^ ions), thus limiting their availability for pathogenic microorganisms in infection sites ^[20]^. Calprotectin coordinates Mn^2+^ through histidine-rich motifs located at the interface of each subunit of the heterodimeric protein. Each subunit also contains a canonical EF-hand motif for binding to calcium ^[21]^. Early biochemical studies on calprotectin indicated that calcium ions are responsible for tuning its Mn^2+^ binding properties, allowing the protein to strongly bind to Mn^2+^ ions only when calcium ions are also available ^[22]^. Based on calprotectin’s unique ability to sequester Mn^2+^ ions and shield from aqueous protons specifically in response to calcium, we reasoned that it should be possible to adapt calprotectin for MRI-based detection of calcium.

Specifically, we hypothesized that in the absence of calcium, a binary mixture of calprotectin and Mn^2+^ ions would effectively shorten the T_1_ and T_2_ relaxation times of water molecules due to relaxation enhancement from free (*i*.*e*., unbound) Mn^2+^. In the presence of calcium, calprotectin would sequester free Mn^2+^ ions, limiting their access to water protons and consequently, increase relaxation times to larger values in inverse proportion to the concentration of free Mn^2+^ ions remaining in solution (**Fig. 1A**). To test our hypothesis, we cloned both subunits of human calprotectin in *E. coli* BL21 cells, substituting the single cysteine residue in each subunit with serine to avoid cross-linking during purification. We purified and reconstituted the 24 kDa heterodimer using metal-affinity chromatography and verified protein function by assaying for calcium-dependent Mn^2+^ binding using a fluorescent dye (**Fig. S1**). Next, we incubated various concentrations of purified calprotectin with Mn^2+^ and measured calcium induced changes in T_1_ and T_2_ relaxation times. These experiments revealed a calcium-dependent increase in relaxation time ranging from 20 ± 4 % to 95 ± 4 % for T_1_and 56 ± 14 % to 201 ± 2 % for T_2_ (*N = 5*) (**Fig. 1B**), which could be respectively visualized as darkening or brightening of MRI signals in standard T_1_ and T_2_ weighted imaging (**Fig. 1C**). Notably, the extent of T_1_ and T_2_ changes obtained with calprotectin are 2 – 6 fold larger than the peak contrast estimated for identical concentrations of the three protein-based MRI sensors (targeting dopamine, serotonin, protein kinase) reported to date (**Fig. S2**) ^[17, 18]^. To further probe the contrast mechanism, we performed relaxometric titrations by treating calprotectin with a range of Mn^2+^ concentrations and measuring T_1_ and T_2_ values in the presence or absence of saturating concentrations of calcium. In calcium-free conditions, we observed a consistent decrease in T_1_ as the amount of Mn^2+^ was increased from 0 to 40 μM (corresponding to 0.25 – 1 molar eq. protein). In contrast, when calcium ions were present, the extent of T_1_ change was significantly smaller (*p < 0*.*01, N = 5*) in the range of 10 – 40 μM Mn^2+^ (**Fig. 2A**). We detected a similar trend in T_2_ values, which decreased with increasing concentrations of Mn^2+^ in the absence of calcium but displayed a substantially smaller change when calcium ions were available (**Fig. 2B**). By fitting the measured T_1_ changes to binding isotherms, we determined that calcium elicits approximately 38-fold increase (*p < 10*^*-5*^, *N = 5*) in calprotectin’s binding affinity for Mn^2+^, consistent with results from previous spectroscopic and calorimetric studies (**Fig. 2C**). To further probe and tune calprotectin’s MRI properties, we introduced point mutations in calprotectin’s Mn^2+^ binding sites, which we predicted from the crystal structure would alter calcium response by modifying Mn^2+^ affinity. We examined the resulting variants using relaxometry (**Fig. S3**) and found one mutant (His_3_Asp → Ala_4_) that displayed lower Mn^2+^ affinity in the calcium-free state relative to wild type calprotectin (*N = 3, p = 0*.*005*), leading to a modest but statistically significant increase in overall calcium response (**Fig. 2C-D**). Finally, we assessed the selectivity of calprotectin’s MRI response towards calcium by incubating a mixture of calprotectin and Mn^2+^ ions with magnesium (50 molar eq.), representing the most abundant divalent cation found in cells. No significant change in T_1_ or T_2_ values could be detected under these conditions (*p ≥ 0*.*2, N = 4*) (**Fig. 2E-F**).

**Figure 1.**
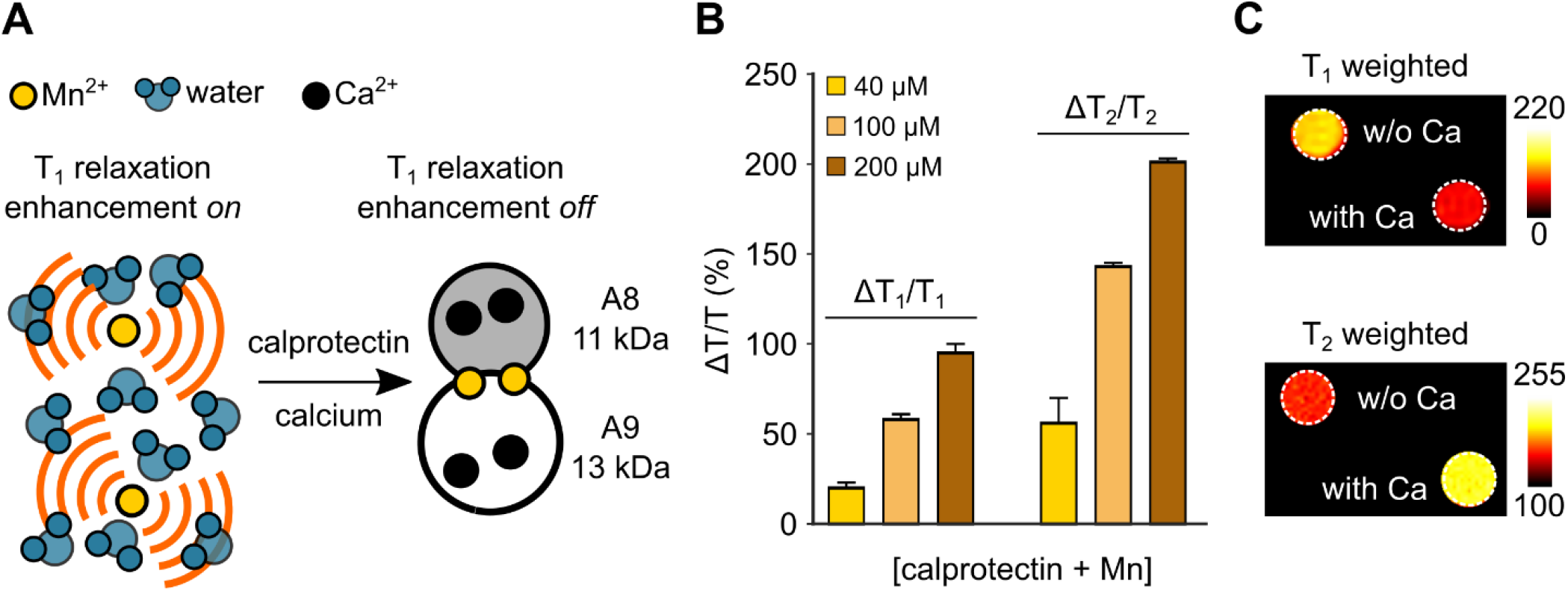
Calcium imaging using a protein-based MRI sensor derived from calprotectin. (A) Proposed mechanism of MRI contrast induced by sequestration of paramagnetic Mn^2+^ions by calprotectin in the presence of calcium. (B) Percent change in T_1_ and T_2_ relaxation times for various concentrations of a binary mixture of calprotectin and Mn^2+^, in response to saturating amounts of calcium. (C) T_1_ and T_2_ weighted images of calcium induced MRI contrast obtained with 200 μM calprotectin. Error bars represent standard error of mean from 5 independent replicates.

**Figure 2.**
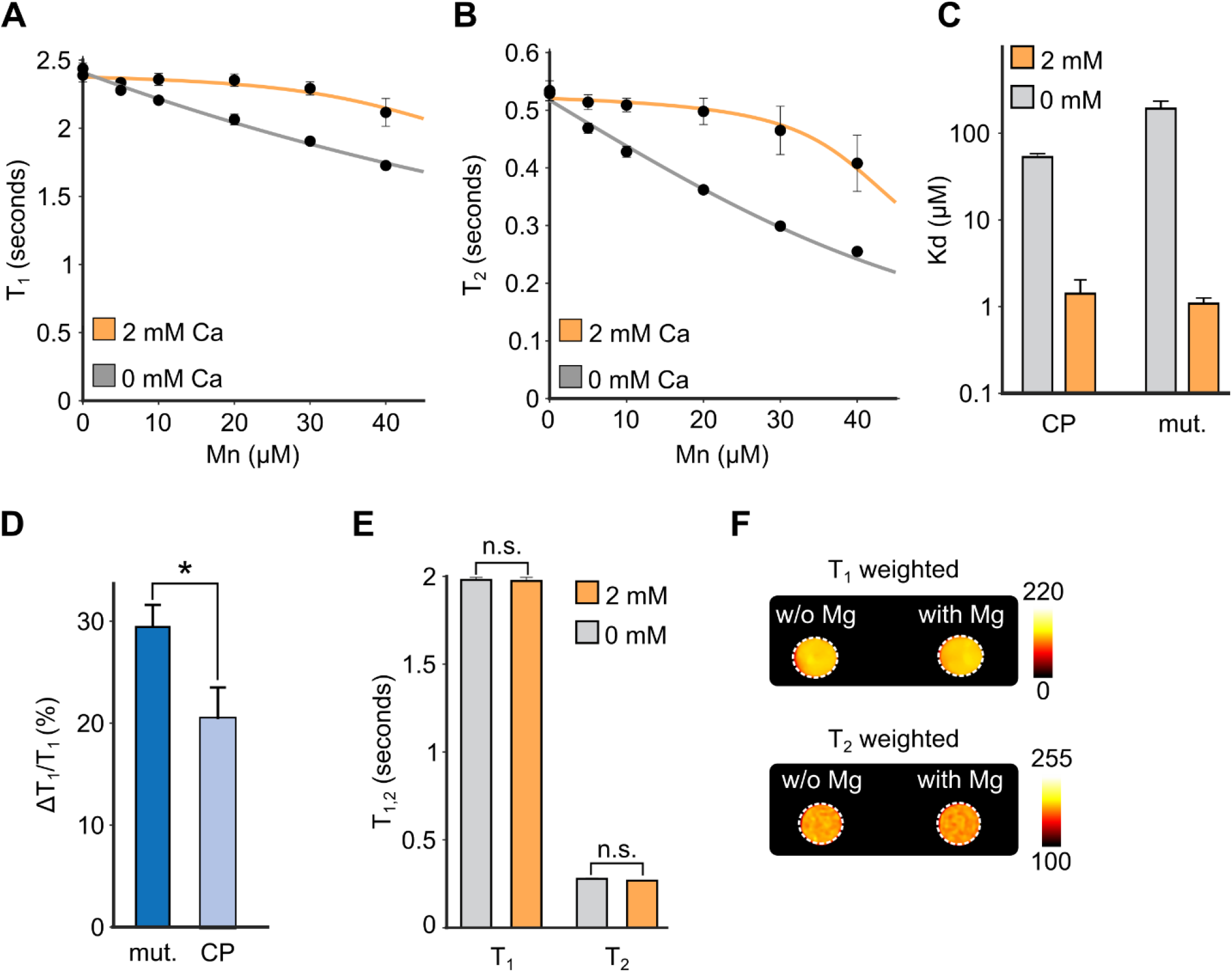
Relaxometric titration of calprotectin-based sensor. In the presence of calcium, (A) T_1_ and (B) T_2_ relaxation times exhibit a significantly smaller decrease with increasing Mn^2+^ concentrations due to Mn^2+^ sequestration by calprotectin. The solid lines represent best fits to equilibrium binding isotherms. (C) Dissociation constants for Mn^2+^ binding to calprotectin (CP) and His_3_Asp variant (mut.) in the presence and absence of saturating calcium, estimated from T_1_ titration results. (D) Comparison of calcium-induced percent change in T_1_ for calprotectin and the His_3_Asp mutant. (E) Calprotectin does not produce a change in T_1_ and T_2_ values or (F) detectable T_1_ and T_2_ weighted contrast in response to saturating amounts of Mg^2+^. Error bars represent standard error of mean from 3 – 5 independent replicates. ^*^ denotes *p < 0*.*05* and n.s. indicates *p > 0*.*05* (Student’s t-test).

Next, we evaluated the dynamic range of calprotectin response by measuring T_1_ and T_2_ changes in buffers consisting of varying concentrations of calcium. For these (and subsequent) experiments, we fixed the amount of calprotectin and Mn^2+^ at 40 and 30 μM, based on Mn^2+^ concentrations that can be readily and safely achieved in cells (**Fig. S4**) as well as the observed Mn^2+^ to protein stoichiometry required for optimum calcium-dependent contrast. In these settings, we detected an 18.7 ± 2.5 % (*p = 0*.*007, N = 3*) increase in T_1_ and a 77.5 ± 1.5 % (*p = 1*.*0 x 10*^*-4*^, *N = 3*) increase in T_2_ over calcium concentrations spanning the full physiological range of (0.1 – 100 μM) (**Figs. 3A, S5**). Our results indicate that calprotectin can be used to detect physiologically relevant calcium concentrations based on an increase in T_1_ and T_2_ relaxation times resulting from calprotectin binding to free paramagnetic Mn^2+^ ions and sequestering them in a low relaxivity state.

**Figure 3.**
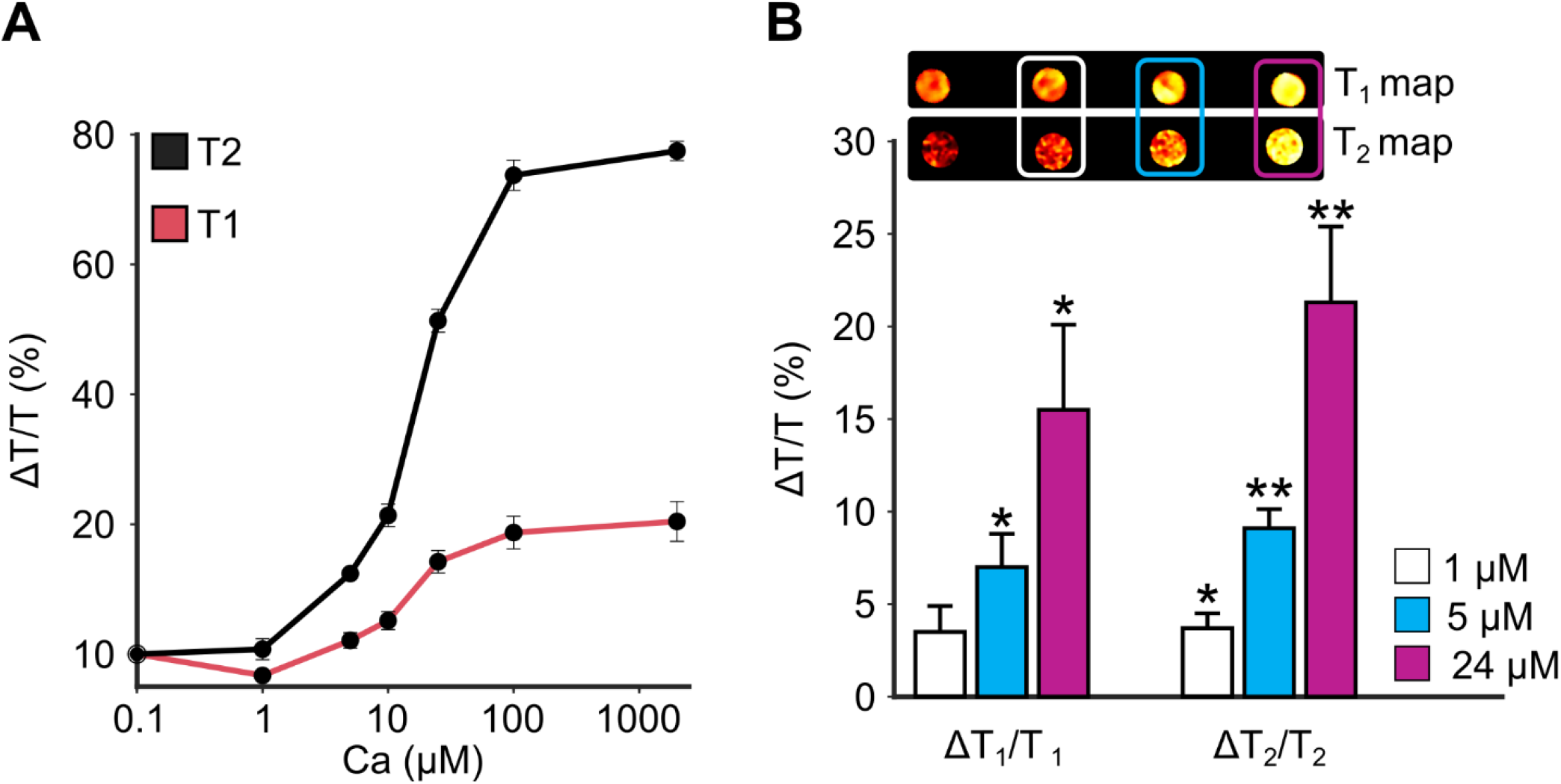
Calcium response properties of calprotectin-based sensor. (A) Percent change in T_1_ and T_2_ relaxation times in response to calcium concentrations spanning the full physiological range. (B) Percent change in relaxation times obtained using calprotectin in a hippocampal cell lysate treated with biologically relevant concentrations of calcium. Calcium response is also represented by voxel-wise mapping of T_1_ and T_2_ values in a representative cell lysate sample. Error bars represent standard error of mean from 4 – 5 independent replicates. ^*^ denotes *p < 0*.*05* and ^**^ indicates *p < 0*.*01* (Student’s t-test).

Finally, as a precursor to genetically encoded imaging in mammalian cells, we examined the sensor’s calcium response in a cellular context using intracellular lysates prepared from a mouse hippocampal cell line (HT-22). Similar lysate preparations have been previously used for *in vitro* validation of protein-based MRI sensors for imaging kinase activity ^[18]^. We supplemented the cell lysate with purified calprotectin and manganese chloride, and measured changes in relaxation times following treatment with calcium concentrations relevant to intracellular signaling. In this cellular milieu, calcium concentrations as low as 5 μM were found to induce a significant increase in T_1_ (7.0 ± 1.8 %, *p = 0*.*02, N = 4*) and T_2_ (9.1 ± 1.0 %, *p = 0*.*0015, N = 4*) (**Fig. 3B**). No change in T_1_ or T_2_ values could be detected when calcium was added to control lysates containing Mn^2+^ but lacking calprotectin (*p > 0. 2, N = 3*) (**Fig. S6**). Importantly, the MRI response obtained with calprotectin is comparable to signal changes reported for existing Gd^3+^ and Mn^3+^ based calcium probes (**Table S1**) and corresponds to amplitudes that can be reliably detected *in vivo* by MRI ^[8, 10, 14]^. These observations indicate that calprotectin’s response properties are appropriate for functional imaging of neural activity in response to various stimulation paradigms, including electrical or optogenetic neuromodulation and chemically induced seizures ^[23]^, which have been shown to elevate resting-state calcium (typically, 50-100 nM in neurons) to tens of micromolar concentrations.

## DISCUSSION

Our findings represent the first *in vitro* proof-of-concept for a protein-based construct for calcium imaging by MRI. The observed changes in T_1_ and T_2_ relaxation times in response to calcium binding are comparable to most synthetic calcium probes and exceed that of previous protein-based neurotransmitter sensors (**Fig. S2**). Furthermore, the ability to image calprotectin response by T_1_ as well as T_2_ weighted MRI may provide an additional advantage in instances where one or the other mechanism may produce contrast that is more clearly distinguishable from background signals due to changes in blood flow, oxygenation, and other physiological effects. Another major advantage, unique to protein-based sensors is the ability to utilize molecular engineering techniques such as directed evolution to readily develop variants with improved functionalities, best illustrated by the prolific expansion of fluorescent calcium indicators starting from parent constructs such as GCaMP ^[15, 24]^. In the future, it may be analogously possible to refine our current sensor by applying classical engineering methods to increase calcium sensitivity, dynamic range, and response amplitude, preliminary proof-of-principle for which is afforded by our mutagenesis studies (**Fig. 2C**). Future efforts may also focus on expressing calprotectin endogenously in neurons and ultimately, *in vivo* by assembling genes for each subunit into a polycistronic construct and packaging in appropriately serotyped viral vectors or using homology independent CRISPR-based techniques to integrate the subunit genes in genomic safe harbor loci ^[25]^. Loading cells with Mn^2+^*in vivo* should be possible using one of several well-established approaches for Mn^2+^ delivery in vertebrates, including dietary supplementation, intraperitoneal injection, and transient opening of the blood brain barrier ^[26]^. If needed, intracellular Mn^2+^delivery may be facilitated through co-expression of divalent metal transporters such as DMT1 ^[19, 27]^. As the first protein-based MRI sensor for calcium, calprotectin represents a key starting point towards engineering genetically targeted constructs for noninvasive, brain-wide imaging of calcium signaling in vertebrates, which has the potential to significantly aid basic research into neural mechanisms involved in learning, memory, addiction, brain injury, and neurodegenerative disorders.

## ACKNOWLEDGEMENTS

We thank members of the Mukherjee lab and our collaborators for helpful discussions. We are grateful to Prof. Elizabeth Nolan (MIT) for introducing us to calprotectins. We thank Prof. Mikhail Shapiro (Caltech) for insightful discussions on MRI and calcium imaging. This work was supported by the California NanoSystems Institute (University of California, Santa Barbara), a NARSAD Young Investigator Award (to AM) from the Brain and Behavior Research Foundation, and a National Institutes of Health R35 Maximizing Investigators’ Research Award (to AM). HFO gratefully acknowledges support from the Errett Fisher Foundation.

## Supporting Information

**Figure S1.**
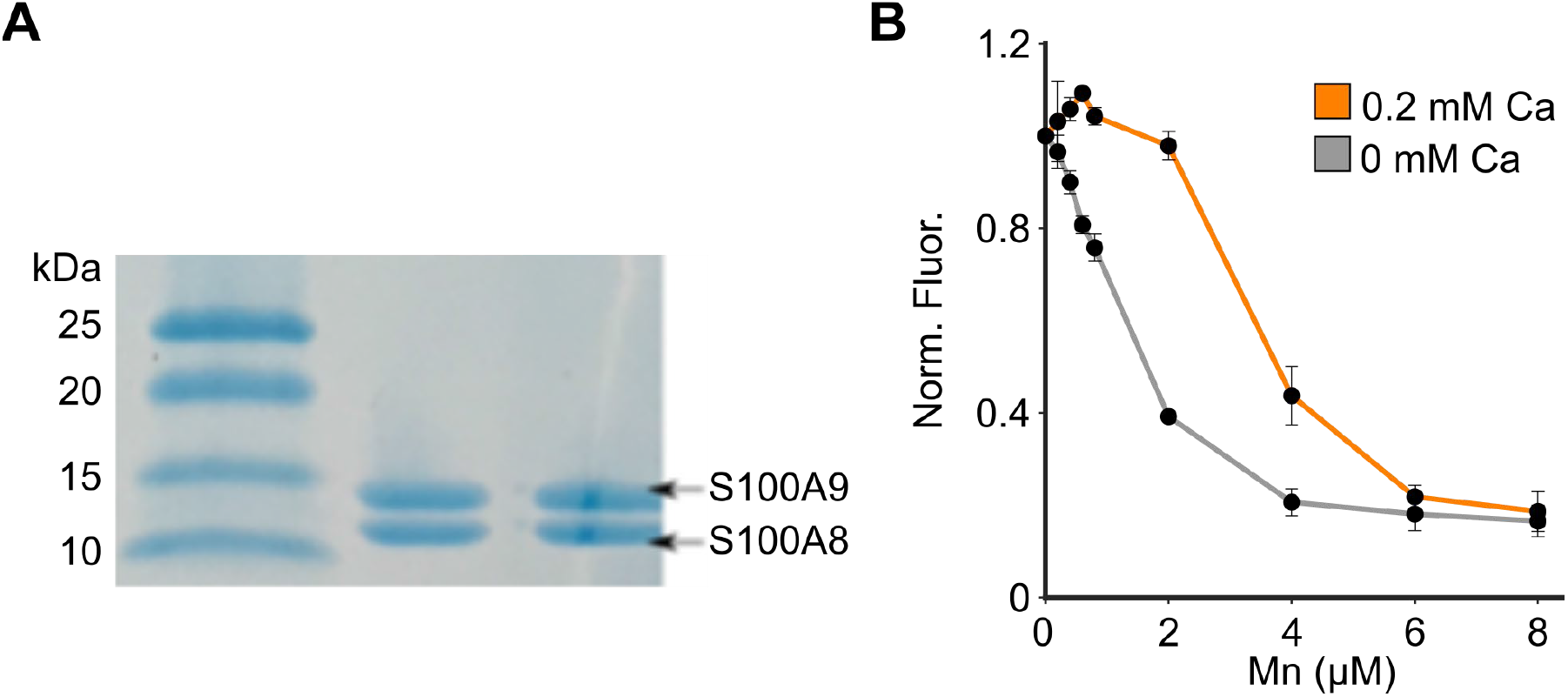
Purification and functional validation of calprotectin. (A) Denaturing gel electrophoresis of reconstituted calprotectin visualized by Coomassie staining. (B) Zinpyr-1 fluorescence^[3]^ titration curve indicating Mn^2+^ binding to calprotectin (4 μM) in the presence of saturating concentrations of calcium.

**Figure S2.**
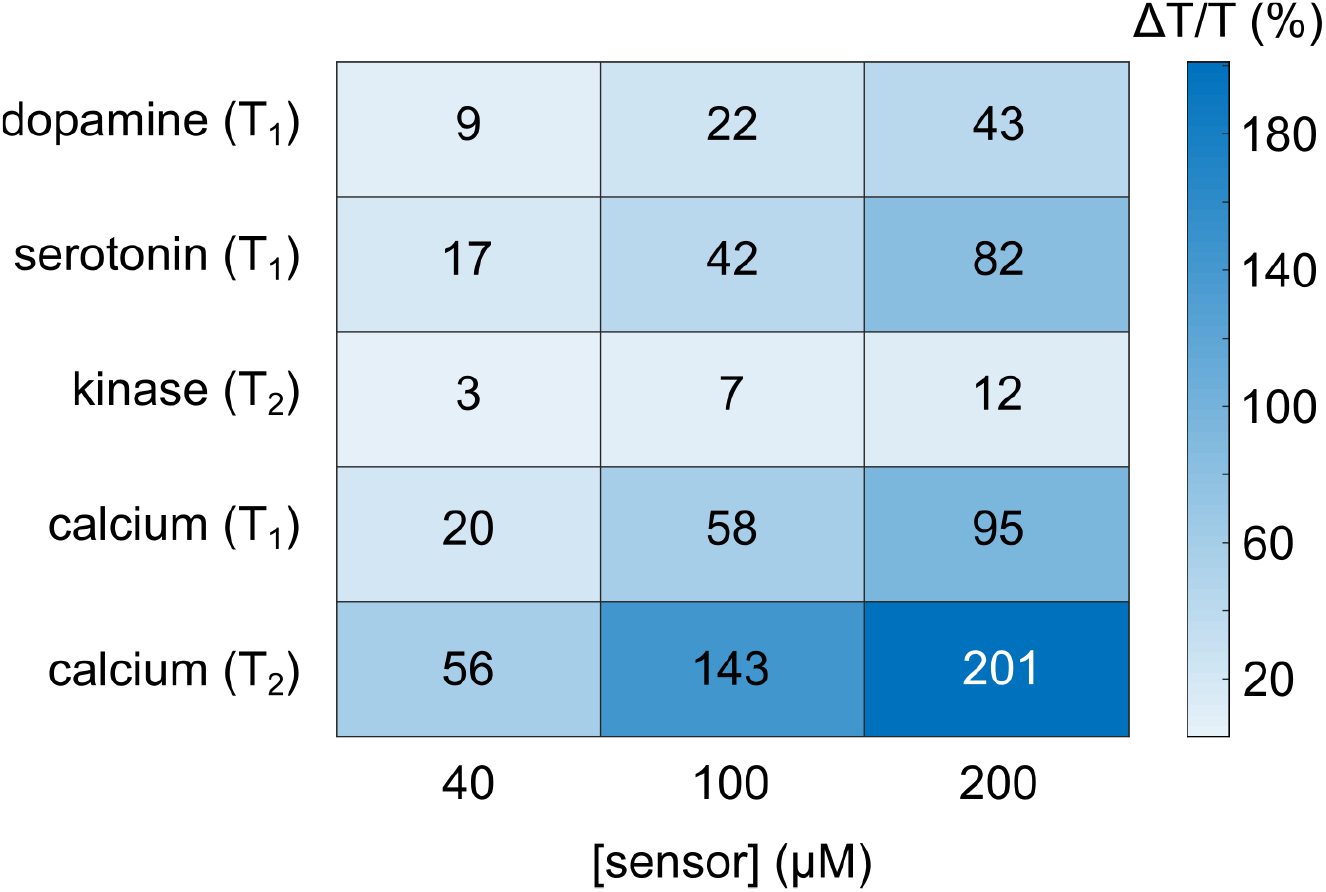
Maximum analyte-induced percent change in relaxation times for protein-based MRI sensors. For the dopamine^[1]^, kinase^[4]^, and serotonin^[6]^ sensors, percent change is calculated based on published relaxivity values

**Figure S3.**
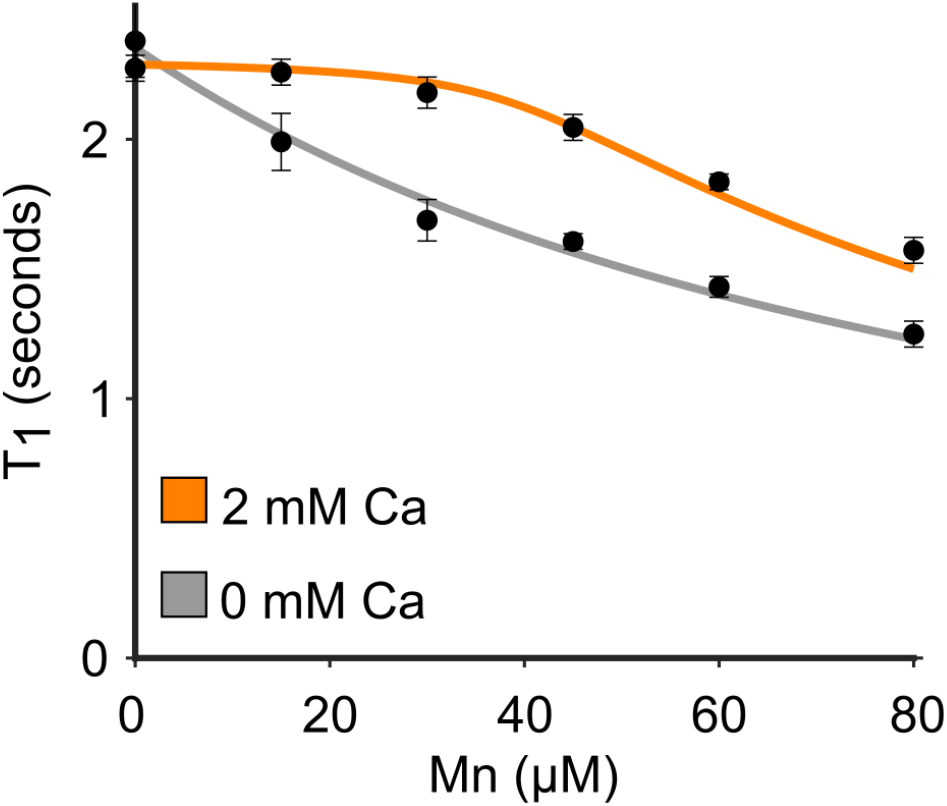
Relaxometric titration of His_3_Asp→Ala_4_ in the presence or absence of calcium.

**Figure S4.**
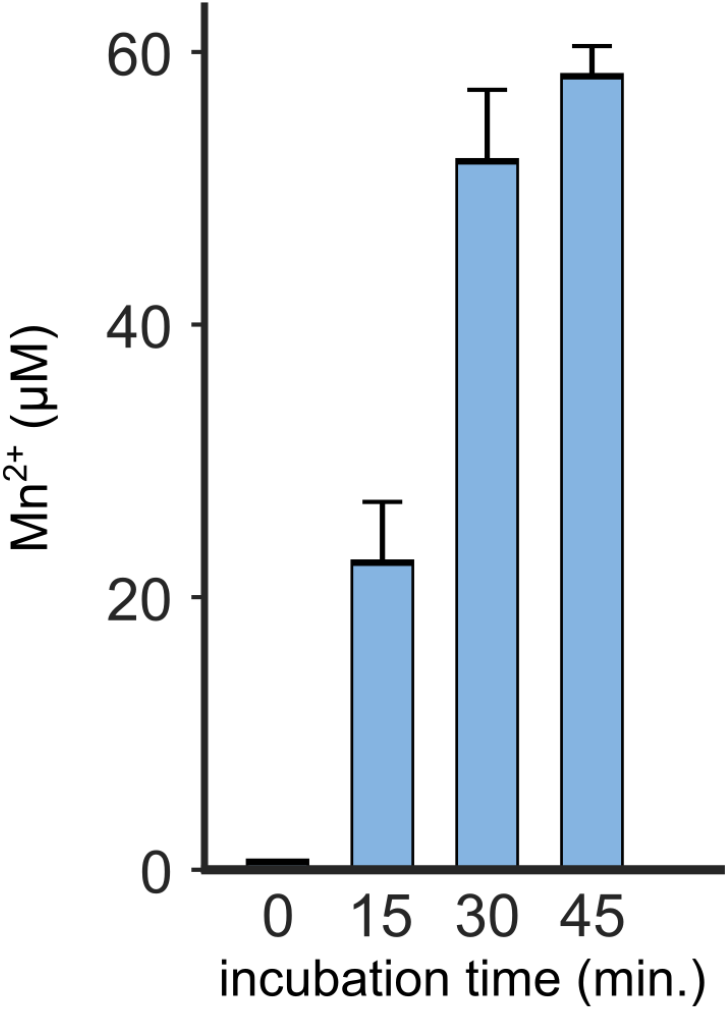
Mn^2+^ uptake in cells. Cells were treated with 100 μM MnCl_2_ for varying durations, rinsed, and analyzed for Mn^2+^ content by ICP/MS.

**Figure S5.**
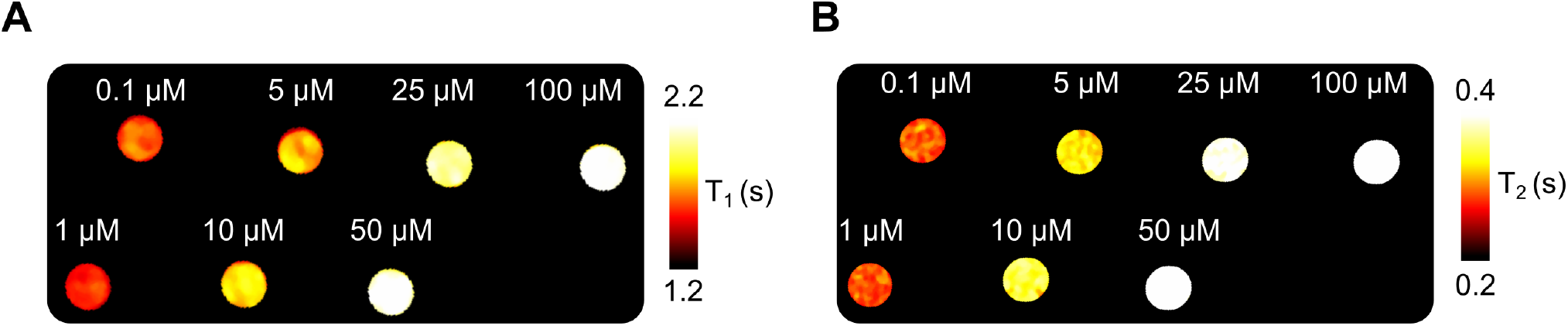
T_1_ and T_2_ response of calprotectin-based calcium sensor. (A) Voxel-wise T_1_ and (B) T_2_ maps of calprotectin response to varying concentrations of calcium.

**Figure S6.**
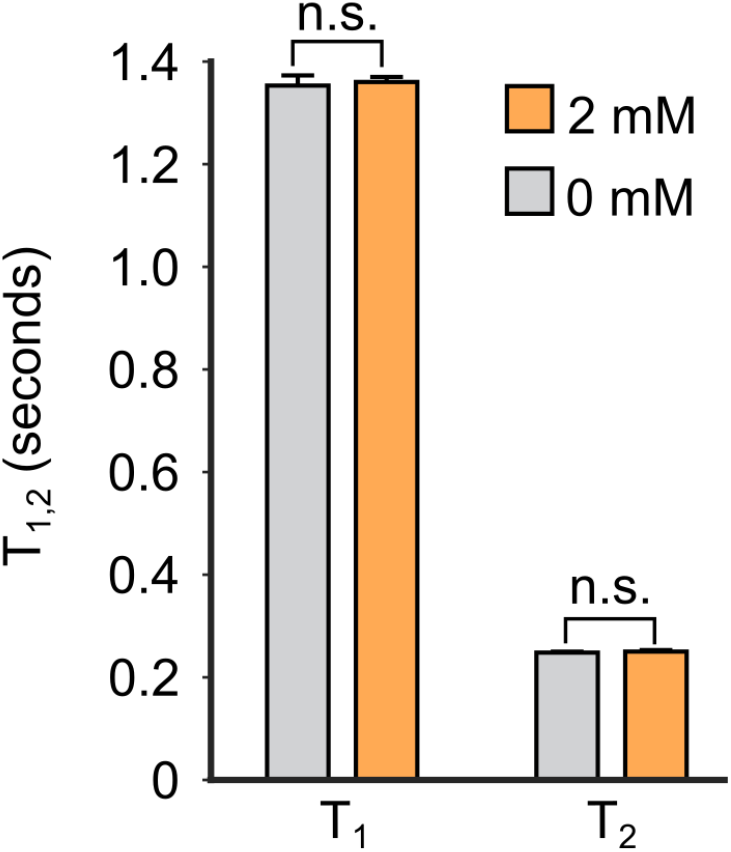
Calcium imaging in HT-22 cell lysate. No T_1_ and T_2_ changes in response to calcium are observed if calprotectin is omitted from the lysate (*i*.*e*., lysate treated with calcium in the presence of Mn^2+^ alone).

**Table S1.**
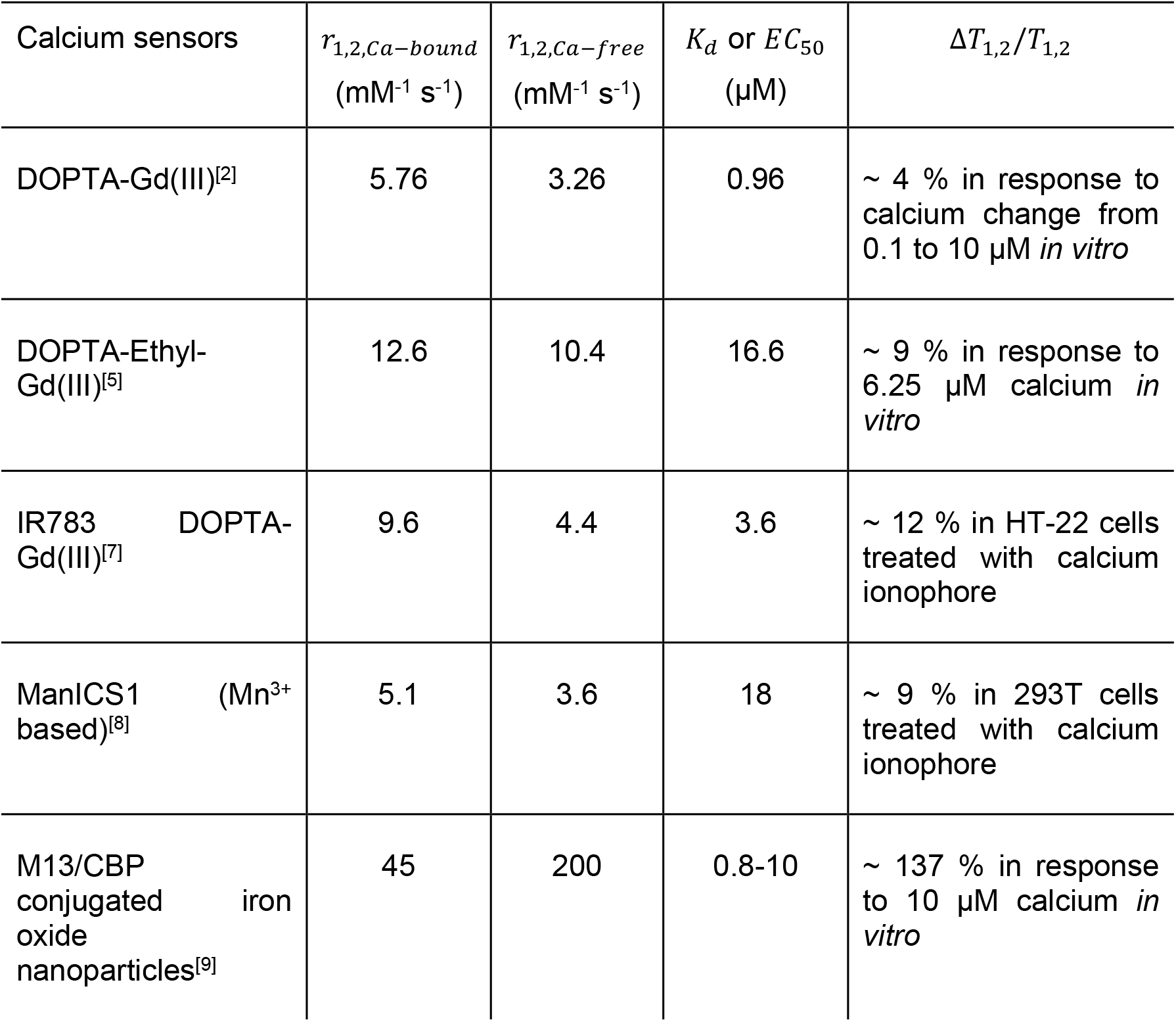
MRI response of intracellular calcium sensors (synthetic)

| Calcium sensors | $r_{1,2,Ca-bound}$<br>(mM<sup>-1</sup> s<sup>-1</sup>) | $r_{1,2,Ca-free}$<br>(mM<sup>-1</sup> s<sup>-1</sup>) | $K_d$ or $EC_{50}$<br>(μM) | $\Delta T_{1,2}/T_{1,2}$ |
| --- | --- | --- | --- | --- |
| DOPTA-Gd(III)<sup>[2]</sup> | 5.76 | 3.26 | 0.96 | ~ 4 % in response to calcium change from 0.1 to 10 μM *in vitro* |
| DOPTA-Ethyl-Gd(III)<sup>[5]</sup> | 12.6 | 10.4 | 16.6 | ~ 9 % in response to 6.25 μM calcium *in vitro* |
| IR783 DOPTA-Gd(III)<sup>[7]</sup> | 9.6 | 4.4 | 3.6 | ~ 12 % in HT-22 cells treated with calcium ionophore |
| ManICS1 (Mn<sup>3+</sup> based)<sup>[8]</sup> | 5.1 | 3.6 | 18 | ~ 9 % in 293T cells treated with calcium ionophore |
| M13/CBP conjugated iron oxide nanoparticles<sup>[9]</sup> | 45 | 200 | 0.8-10 | ~ 137 % in response to 10 μM calcium *in vitro* |

## SUPPLEMENTAL METHODS

### Materials and reagents

Zinpyr-1, a fluorescent turn-off dye for Mn^2+^ was purchased from Adipogen Corporation (San Diego, CA, USA). HT-22 mouse hippocampal cell lines were purchased from Thermo Fisher Scientific (Waltham, MA, USA). Reagents for Gibson assembly and mutagenesis were purchased from New England Biolabs (Ipswich, MA, USA). All other chemicals and reagents were purchased from or MilliporeSigma (St. Louis, MO, USA) or Thermo Fisher Scientific unless otherwise specified.

### Cloning and mutagenesis

*E. coli* glycerol stocks containing *H. sapiens* ORF clones for each calprotectin subunit – S100A8 (RefSeq Accession: NM_001319201.1) and S100A9 (RefSeq Accession: NM_002965.3) – were purchased from GeneCopoeia (Rockville, MD, USA). Following isolation of the ORF plasmids, S100A8 and S100A9 genes were amplified by PCR and sub-cloned by Gibson assembly into a pET26b expression vector, which appended an N terminus hexahistidine tag to each subunit. Point mutations were introduced by site directed mutagenesis to generate the following single mutants: S100A8 C50S, S100A9 C11S, and the triple mutants: S100A8 C50S/H91A/H95A, S100A9 C11S/H28A/D38A. All gene sequences were verified by Sanger sequencing (Genewiz, South Plainfield, NJ, USA) prior to transforming the resulting plasmids in *E. coli* BL21(DE3) cells for protein expression.

### Protein purification

Each subunit of calprotectin (harboring a Cys→Ser substitution to prevent cross-linking^[10]^) was expressed individually in *E. coli* cells. For protein expression, 5 ml of overnight culture was inoculated in 500 mL LB broth supplemented with 50 μg/ml kanamycin. Cells were grown at 37 °C until they reached an OD_600_ of 0.6 at which point protein expression was induced by adding 0.5 mM isopropyl-β-D-1-thiogalactopyranoside (IPTG). Protein expression was continued for another 3 – 4 h before collecting cells by centrifugation at 4000 x g for 8 minutes at 4 °C. Cell pellets corresponding to each calprotectin subunit were combined by resuspending in 15 ml lysis buffer (50 mM Tris hydrochloride, 100 mM NaCl, 1 mM EDTA, 0.5% Triton-X, 1 mM phenylmethylsulfonyl fluoride, pH 8) and sonicated using a QSonica Q500 cell disruptor with the following parameters – pulse length, 2.5 min; pulse frequency: 30 s on/10 s off, and pulse amplitude: 20 – 40%. Following sonication, the solution was centrifuged at 30,000 x g for 20 minutes at 4 °C. The supernatant was discarded, and the resulting pellet was resuspended in 30 ml lysis buffer. Sonication and centrifugation steps were repeated two more times before gently solubilizing the final pellet in 60 ml solubilization buffer (50 mM Tris hydrochloride, 100 mM NaCl, 4 M guanidine hydrochloride, pH 8). The solubilized pellet was sonicated and centrifuged one final time before exchanging into buffer containing 20 mM HEPES (pH 8.0) and 10 mM imidazole, by at least 5 rounds of centrifugation using Amicon Ultra centrifugal filter units (MilliporeSigma, 3 kDa MWCO). The retentate was purified using nickel affinity chromatography by loading on a HisTrap column (GE Healthcare, Piscataway, NJ, USA) and eluting with a linear gradient of 20-500 mM imidazole in 20 mM HEPES buffer (pH 8.0). Eluted fractions were analyzed by gel electrophoresis and protein fractions corresponding to the two expected bands (11.8 kDa for His_6_ tagged S100A8 C50S and 14.2 kDa for His_6_ tagged S100A9 C11S) were pooled and exchanged by centrifugal filtration into 20 mM HEPES buffer (pH 7.5). Concentration of the final purified protein was estimated using the Bradford assay (Biorad, Hercules, CA, USA).

### Fluorescence titration assay of Mn^2+^ binding

Stock solutions of MnCl_2_ (1 mM) and CaCl_2_ (40 mM) were prepared by dissolving the respective hydrated salts in deionized water. Stock solution of Zinpyr-1 (1 mM) was prepared by dissolving in dimethyl sulfoxide and stored in light– tight conditions. Binding assays were conducted in 300 μL reaction volumes comprising 1 μM Zinpyr-1, 4 μM calprotectin, 0 or 200 μM CaCl_2_, and various Mn^2+^ concentrations in the 0 – 8 μM range^[10]^. The resulting mixture was allowed to incubate briefly at room temperature before acquiring fluorescence measurements on a Tecan Spark microplate reader, by exciting at 450 nm and scanning emission from 485 to 630 nm. The monochromator slit width and gain were respectively set at 20 nm and 70.

### Cell culture, lysate preparation, and intracellular Mn^2+^ measurements

Adherent mouse hippocampal HT-22 cell lines were grown in Dulbecco’s Modified Eagle Media supplemented with 10 % fetal bovine serum, 100 U/mL penicillin, and 100 μg/mL streptomycin, at 37 °C, 5 % CO_2_ in a humidified chamber. Once the cells reached 70 – 80 % confluency, they were lysed using the RIPA Lysis Buffer System (Santa Cruz Biotechnology, Dallas, TX, USA) according to the manufacturer’s instructions. The lysate was briefly spin filtered using Amicon Ultra centrifugal filter units to remove the lysis reagents and reconstituted to at least 60% (v/v) in HEPES buffer (pH 7.4). For measuring Mn^2+^ uptake, cells were cultured in 6-well plates, supplemented with 100 μM MnCl_2_ for different durations, rinsed with sterile phosphate buffered saline, lysed using RIPA buffer and assayed for Mn^2+^ content using an inductively coupled plasma – atomic emission spectroscopy (Perkin Elmer 5300 DV). ICP-grade Mn^2+^ was used as a calibration standard.

### *In vitro* MRI

For all MRI experiments, known concentrations of calprotectin and Mn^2+^ were first added to 20 mM HEPES buffer (pH 7.4) or HT-22 cell lysate in 200 μL tubes and treated with various amounts of calcium (or magnesium to test for specificity). To minimize susceptibility differences during MRI, the tubes were placed in water-filled (1 % w/v) agarose molds housed inside a custom-built 3D printed phantom. All MRI experiments were conducted in a Bruker 7 T vertical bore scanner using a 66 mm diameter transceiver coil. T_1_ scans were acquired using a rapid acquisition with relaxation enhancement (RARE) spin echo sequence with the following parameters: echo time, T_E_: 12.6 ms, RARE factor: 4, field of view: 4.7 cm x 4.7 cm, matrix size: 128 x 128, slice thickness: 1.5 – 2 mm, and variable repetition times, T_R_: 173.5, 305.9, 458.6, 638.8, 858,7, 1141.1, 1536.1, 2198.4, and 5000 ms. T_2_ scans were acquired using a Carr Purcell Meiboom Gill (CPMG) sequence with the following parameters: T_R_: 2 s, field of view: 4.7 cm x 4.7 cm, matrix size: 256 x 256, slice thickness: 1.5 – 2 mm, and multiple echo times, T_E_: 20.9, 31.4, 41.9, 52.3, 62.8, 73.3, 83.7, 94.2, and 104.7 ms.

### MR image analysis

MRI signal intensities were determined by drawing regions of interest on axial sections of the sample tubes and estimating mean voxel intensity using ImageJ (NIH). T_1_ values were determined by fitting the resulting signal intensities (*SS*) to the equation: *SS/SS*=1 – *eeeeee* (–*TT/TT*), which describes the growth of longitudinal magnetization by T_1_ relaxation following a 90 degree excitation pulse. Likewise, T_2_ values were estimated by fitting signal intensities corresponding to the last 9 echoes, to the equation: *SS/SS*= *eeeeee* (–*TT/TT*), which describes the decay of transverse magnetization by T_2_ relaxation. T_1_ and T_2_ weighted images were acquired using RARE and CPMG pulse sequences as above but at a single repetition (T_R_= 459 ms) or echo time (T_E_ = 94 ms). For the calcium titration results, MR images are depicted as T_1_ and T_2_ maps to facilitate visualization of calcium-induced contrast. These maps were generated by fitting the MRI signal intensity in each voxel (within a region of interest) to the aforementioned exponential growth and decay equations. The resulting images were smoothed using a 2 – 4 pixel wide median filter and displayed on a linear 8-bit color scale. Least squares regression and voxel-wise mapping were implemented using Matlab (v. 2018a).

### Analysis of Mn^2+^ binding

Mn^2+^ binding to calprotectin was assumed to proceed by simple bimolecular interaction as follows: CP + Mn^2+^ ↔ CP.Mn. A dynamic mass balance model for this reaction can be constructed as follows:

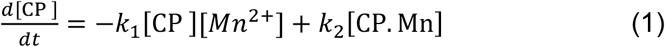

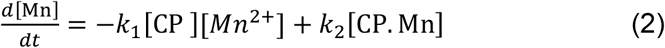

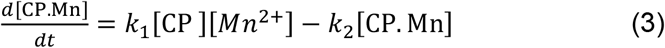

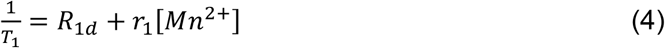

Here, *k*_1_and *k*_2_represent on and off rates for complex formation. The equilibrium dissociation constant can be calculated as *K*_*d*_= *k*_2_*/k*_1_. *T*_1_relaxation time was calculated based on the amount of free Mn^2+^ remaining in solution at equilibrium and experimentally measured value of *r*_1_(5.6 mM^-1^ s^-1^) for pure MnCl_2_ at 7 T. *R*_1d_represents diamagnetic T_1_ relaxation rates for aqueous buffer or cell lysate, typically around 0.4 s^-1^. Two sets of five independent replicates of *T*_1_measurements were fit to the above equations to estimate two *K*_*d*_values – corresponding to Mn^2+^ affinity in the calcium-free state and calcium saturated conditions. A similar approach could also be used derive *KK*values based on T_2_ relaxivity results. However, the T_2_ effect of protein-bound Mn^2+^ is likely to be non-negligible (and more significant than T_1_ effects) on account of outer sphere contributions, which should ideally be accounted for in the model by including *r*_2_values for the CP.Mn complex. As a result, we based our *K*_*d*_estimates on experimental T_1_ data. Some of the prior studies on Mn^2+^ binding to calprotectin (using EPR spectroscopy) has also involved 2-site binding models to account for contributions from a second Mn^2+^ binding site^[10]^. Non-specific Mn^2+^ binding has also been suggested to occur in the EF hand motifs, particularly when unoccupied by calcium^[11]^. Incorporating these additional contributions did not lead to acceptable fits or parameter estimates based on our experimental data. Nonlinear least squares regression was implemented using Matlab (v. 2018a).

### Data analysis

All measurements are reported as mean ± standard error based on experiments conducted using 3 – 5 independent biological replicates. For linear least squares regression, quality of model fits was determined from the regression coefficient. For nonlinear least squares regression, fit quality was judged by inspecting residuals. Pairwise comparisons were performed using the Student’s t-test with significance level set at 0.05.

### Theoretical estimation of MRI signal change

To compare the maximum amplitude of calcium response observed in calprotectin with the three previously published protein-based MRI constructs (for sensing dopamine^[1]^, serotonin^[6]^, and kinase^[4]^), we calculated the theoretical T_1_ or T_2_ change obtained using identical concentrations of the latter sensors (40 – 200 μM) under saturating ligand conditions. Our calculations assumed a background relaxation rate of 0.4 s^-1^ (T_1_) or 1.9 s^-1^ (T_2_) based on experimental measurements acquired using our 7 T vertical bore Bruker MRI. Relaxivity values in ligand-free and ligand-bound (or aggregated) states of the sensor proteins were based on published values, which were acquired at a slightly lower field strength (4.7 T).

### DNA sequences of genetic constructs engineered in this work

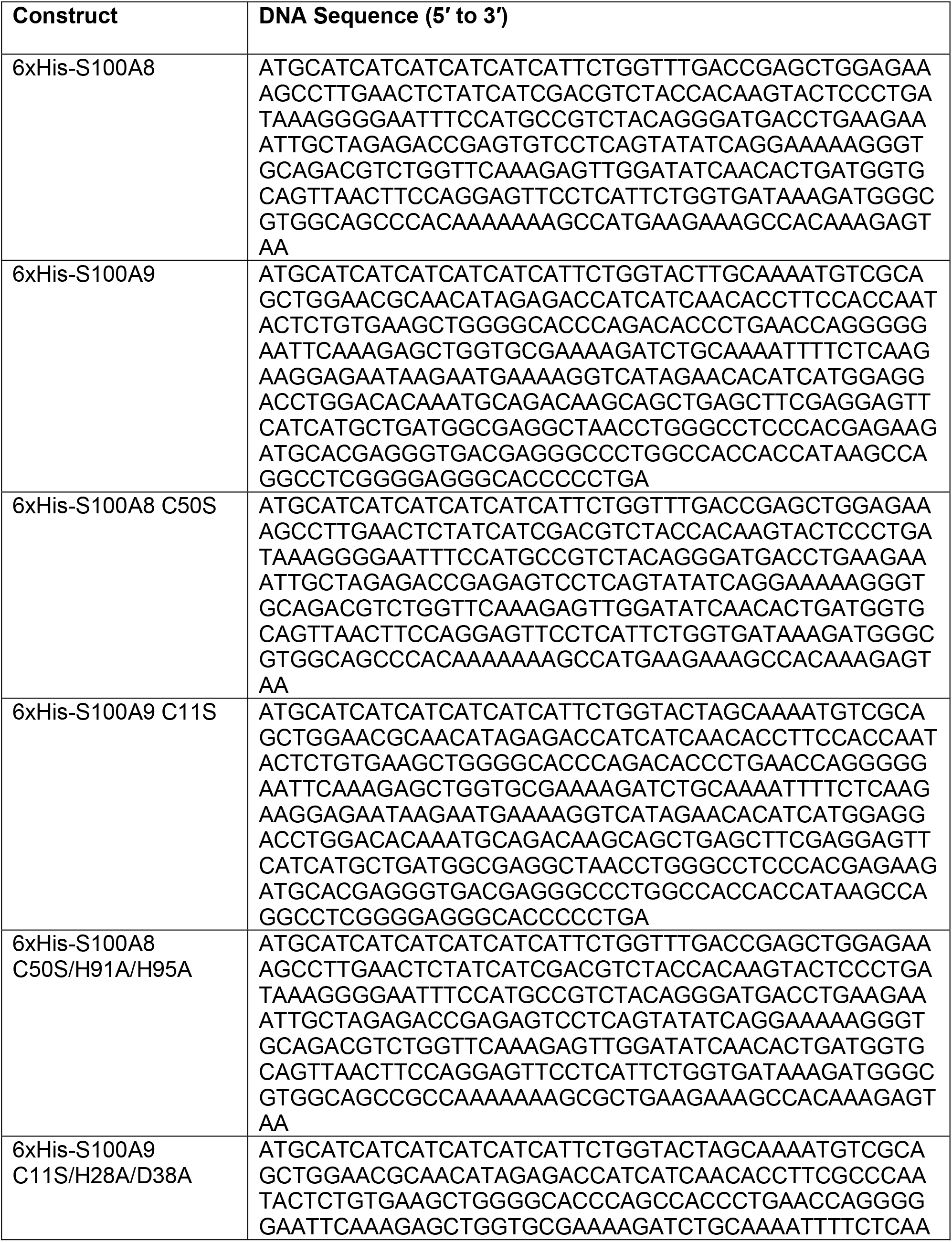
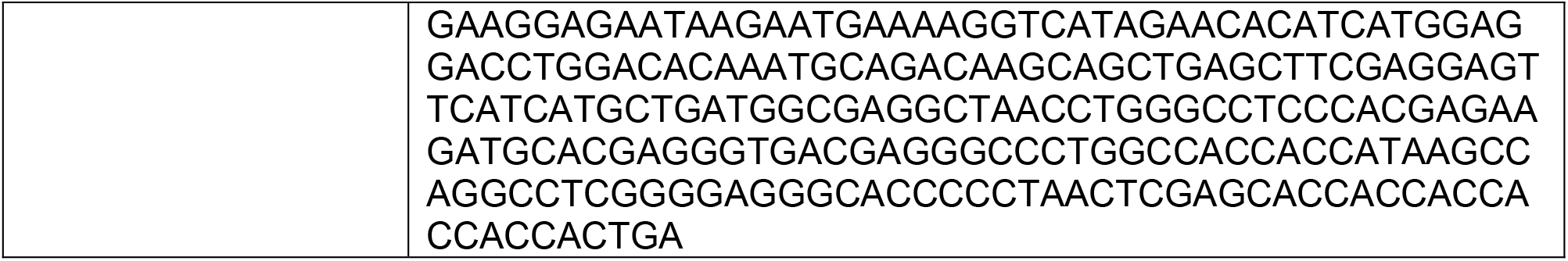

